# Thyroglobulin Interactome Profiling Uncovers Molecular Mechanisms of Thyroid Dyshormonogenesis

**DOI:** 10.1101/2020.04.08.032482

**Authors:** Madison T. Wright, Logan Kouba, Lars Plate

**Affiliations:** Department of Chemistry, Vanderbilt University, Nashville, TN; Department of Biological Sciences, Vanderbilt University, Nashville, TN

**Keywords:** affinity purification–mass spectrometry, congenital hypothyroidism, interactomics, proteostasis, thyroglobulin

## Abstract

Thyroglobulin (Tg) is a secreted iodoglycoprotein serving as the precursor for T3 and T4 hormones. Many characterized Tg gene mutations produce secretion-defective variants resulting in congenital hypothyroidism (CH). Tg processing and secretion is controlled by extensive interactions with chaperone, trafficking, and degradation factors comprising the secretory proteostasis network. While dependencies on individual proteostasis network components are known, the integration of proteostasis pathways mediating Tg protein quality control and the molecular basis of mutant Tg misprocessing remain poorly understood. We employ a multiplexed quantitative affinity purification–mass spectrometry approach to define the Tg proteostasis interactome and changes between WT and several CH-variants. Mutant Tg processing is associated with common imbalances in proteostasis engagement including increased chaperoning, oxidative folding, and routing towards ER-associated degradation components, yet variants are inefficiently degraded. Furthermore, we reveal mutation-specific changes in engagement with N-glycosylation components, suggesting distinct requirements for one Tg variant on dual engagement of both oligosaccharyltransferase complex isoforms for degradation. Modulating dysregulated proteostasis components and pathways may serve as a therapeutic strategy to restore Tg secretion and thyroid hormone biosynthesis.

## INTRODUCTION

Thyroid hormone biosynthesis is an intricate and multifaceted process involving a sequence of biochemical reactions (Carvalho & Dupuy, 2017; Dai *et al*, 1996; Fayadat *et al*, 1999; Di Jeso & Arvan, 2016). Triiodothyronine (T3) and thyroxine (T4) hormones are necessary for normal growth and development in utero and early childhood, and go on to regulate primary metabolism in adulthood (Citterio *et al*, 2019; Oetting & Yen, 2007). Hypothyroidism and dyshormonogenesis stemming from mutations or damage to the biosynthetic components ultimately results in decreased or complete loss in production of T3 and T4. Congenital hypothyroidism (CH) affects approximately 1:2000 to 1:4000 newborns, and if not detected and addressed can lead to severe and permanent neurological damage, including mental retardation (Chaker *et al*, 2017; Rose *et al*, 2006). A critical gene involved in thyroid hormone biogenesis and CH pathology is thyroglobulin (Tg) encoding the prohormone precursor protein for T3 and T4 (Fig 1A). There are 176 documented Tg mutations that impair proper production, folding, or processing leading to dyshormonogenesis (Citterio *et al*, 2019). Missense mutations resulting in full-length but folding-incompetent Tg disrupt normal protein homeostasis (proteostasis) and lead to decreased or complete loss of Tg protein secretion into the thyroid follicular lumen, a key step in hormone production. Instead, mutant Tg variants accumulate within the endoplasmic reticulum (ER) of thyroid follicular cells and are ultimately degraded (Kim *et al*, 1996).

**Figure 1.**
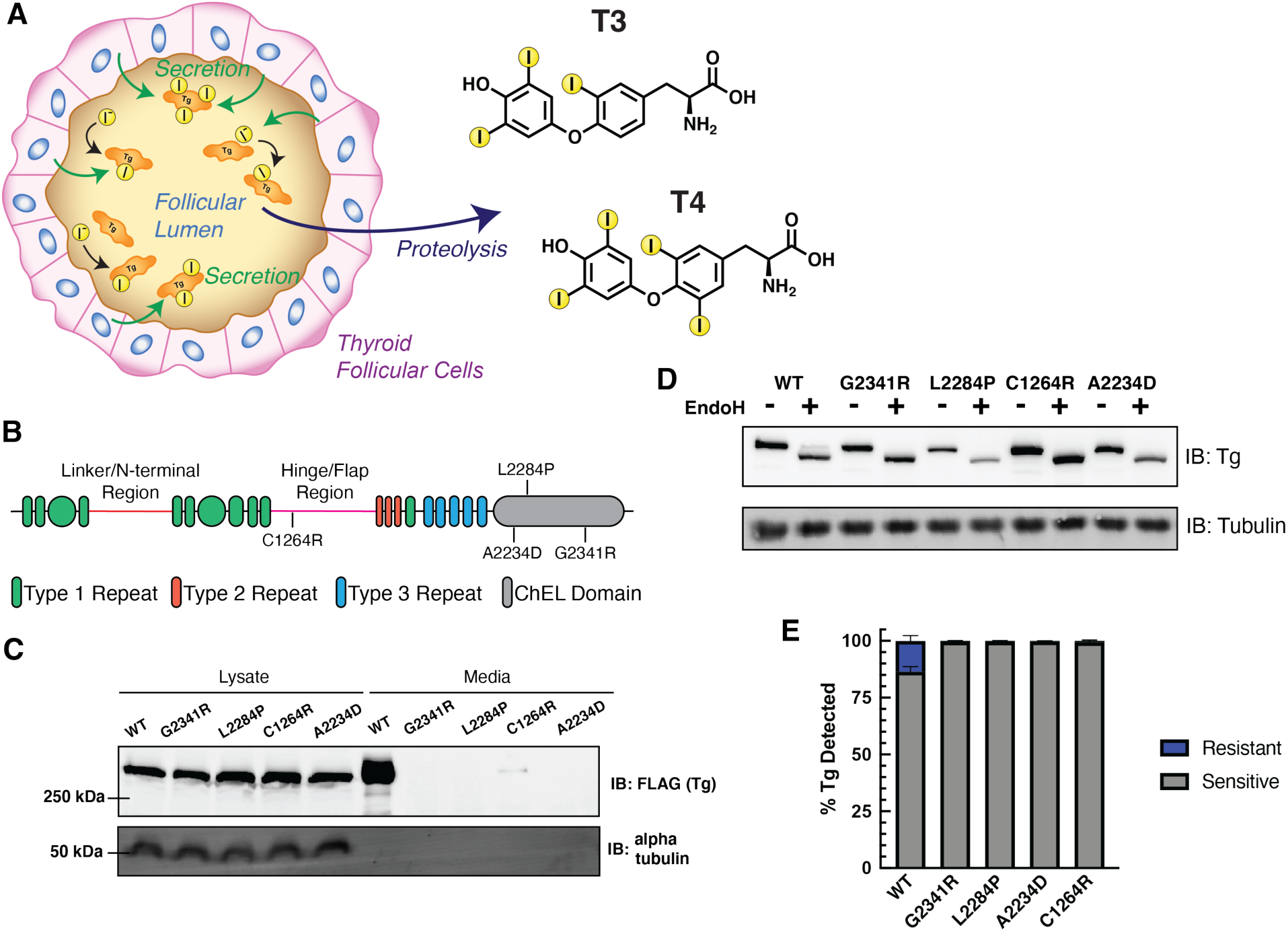
Distinct Tg mutants present secretion defects. **A**. Schematic detailing Tg processing and subsequent hormone production. Tg is synthesized in follicular cells and secreted into the follicular lumen where it undergoes iodination and is stored. Tg is later taken up and proteolyzed leading to the liberation of T3 and T4 hormones. **B**. Schematic of Tg domain organization consisting of cysteine rich repeats, a linker/N-terminal region, and hinge/flap region followed by a cholinesterase like (ChEL) C-terminal domain. **C**. Immunoblot for Flag-tagged Tg expressed in transiently transfected HEK293T cells. All Tg variants are detected in the lysate while only WT is detected in cell culture media. **D**. Western blot for Tg probing EndoH sensitivity to remove high-mannose glycans of ER localized Tg. **E**. Quantification of EndoH sensitivity in D. All Tg mutants are 100% EndoH sensitive, showing they are retained within the ER and model a hypothyroidism phenotype. Error bars show SEM for 4 biological replicates.

While many CH-associated folding-incompetent Tg mutations have been documented, the molecular mechanisms of Tg folding and processing controlled by the proteostasis network (PN), consisting of chaperones, co-chaperones, folding enzymes, trafficking factors, and degradation factors, remain incompletely understood. Coordination of these PN components ensures the proper folding, trafficking, and degradation of clients such as Tg through a process cumulatively referred to as protein quality control (PQC) (Hartl *et al*, 2011; Balchin *et al*, 2016; Sun & Brodsky, 2019). Previous studies have shown that CH-associated Tg mutants exhibit increased interactions with individual PN components including BiP, GRP94, PDIA3, CANX, and CALR, that aid in folding and processing (Menon *et al*, 2007; Baryshev *et al*, 2004; Park & Arvan, 2004; Hishinuma, 1999; Muresan & Arvan, 1997; Di Jeso *et al*, 2005; Kim & Arvan, 1993). Nonetheless, it remains unclear which of these components are responsible for the improper processing of mutant Tg. The current collection of known interactors, identified through traditional immunoprecipitation and immunoblotting strategies, is likely limited as these methods are not conducive to discovery-based investigations. Additionally, little work has focused on characterizing mutation specific changes in PN engagement. Identifying the complete Tg interactome and defining the molecular mechanisms of altered PN engagement for mutant Tg variants may reveal areas of PQC that can be targeted therapeutically to rescue the secretion of these CH-associated variants. No disease modifying therapies currently exist to restore secretion of destabilized Tg, but devising such strategies would be particularly critical considering the increased prevalence of dyshormonogenesis amongst newborns and complications arising from the current “gold standard” of hormone therapy treatments in the clinic (Olivieri *et al*, 2015; Chaker *et al*, 2017). Modulation of individual PN components or entire pathways has shown recent promise as a potential therapeutic strategy to combat a number of protein folding diseases, including light-chain amyloidosis (AL), transthyretin (TTR) amyloidosis, and polyglutamine (polyQ) associated neuropathies (Plate & Wiseman, 2017; Hetz *et al*, 2019; Cooley *et al*, 2014; Chen *et al*, 2014). Such therapies could be similarly effective at restoring Tg secretion and subsequent hormone biosynthesis.

Here, we present a quantitative interactome proteomics method that allowed us to globally profile several CH-associated mutant Tg variants. Unlike previous proteostasis interactome studies (Pankow *et al*, 2015; Plate *et al*, 2019; Doan *et al*, 2019), the multiplexing capabilities used here enable a head-to-head comparative analysis of five distinct protein variants at once. While chaperone complexes and client recognition for select chaperones have been mapped (Taipale *et al*, 2014; Behnke *et al*, 2016; Christianson *et al*, 2012), system-wide investigations into PN processing of individual client proteins are lacking. The current study describes the identification of a comprehensive PN interactome for WT Tg and several secretion deficient mutant variants. Comparison of the PN interactome for the CH-associated mutant variants to WT Tg allowed us to gain mechanistic insights into shared protein quality control defects that are responsible for the loss of secretion of all destabilized variants. Our data supports a model whereby the destabilized Tg variants are retained intracellularly through increased interactions with chaperoning and oxidative protein folding pathway components. We also find evidence that Tg mutants are increasingly routed towards ER-associated degradation (ERAD), but degradation cannot be completed due to failures in retrotranslocation, retention by ER chaperone networks, or overall lower engagement of proteasomal degradation machinery. At the same time, we find mutation specific interactome remodeling with components of N-glycosylation, downstream glycan processing, and lectin-assisted protein folding components. Mutant-specific interaction changes suggest that such Tg variants have distinct imbalances associated with their aberrant folding and processing within the ER, leading to the loss of secretion.

## RESULTS

### Distinct Thyroglobulin Mutants Present Common Secretion Defects

Tg is a large 330 kDa multidomain protein consisting of extensive cysteine-rich repeat regions and a C-terminal cholinesterase like domain (ChEL) (Fig. 1B & Fig. S1A) (Coscia *et al*, 2020). We focused on a set of single-point mutations that lead to impaired Tg secretion in human CH patients (A2234D and C1264R) and in a mouse model of thyroid hormone deficiency and goiter (L2284P) (Kim *et al*, 1998; Caputo *et al*, 2007; Hishinuma, 1999). A2234D and L2284P occur in the ChEL domain, which serves as an intramolecular chaperone playing a critical role in Tg folding, dimerization, and secretion (Lee *et al*, 2008, 2009). Our analysis also included a previously uncharacterized ChEL mutation at a conserved glycine (G2341R), which is located adjacent to L2284 and A2234. We contrasted the ChEL mutations to the C1264R variant in the hinge/flap region (Fig. S1A-D).

We transiently transfected HEK293T cells with FLAG-tagged expression constructs of either WT or the respective mutant Tg variants. We detected all Tg variants at similar levels in lysate samples, while only WT Tg was detected in the media, confirming the secretion defect of CH mutations (Fig. 1C). C1264R Tg was occasionally detected at trace amounts in the media indicating low residual secretion (< 1-2% of WT). These results are in accordance with previous Tg studies (Pardo *et al*, 2009; Lee *et al*, 2011; Hishinuma, 1999; Kim *et al*, 1998). WT Tg undergoes extensive glycosylation within the ER prior to being trafficked and further modified in the Golgi apparatus, while the folding-incompetent CH-associated mutations are trapped within the ER preventing Golgi modifications. To investigate Tg localization and glycosylation state we performed EndoH digestions on the transfected HEK293T lysates. All mutants were EndoH sensitive, indicating they had not traversed the medial Golgi apparatus as EndoH specifically cleaves ER associated high-mannose glycans. In contrast, WT Tg separated into two distinct EndoH resistant and EndoH sensitive populations indicating that WT Tg was able to fold within the ER and traverse the secretory pathway (Fig. 1D-E). These data further confirm that the C-terminal FLAG epitope tag does not influence Tg glycosylation. Overall, our results confirm that FLAG epitope tagged Tg constructs do not show altered processing and serve as a useful model system to probe PQC dynamics for WT and CH-associated, secretion-deficient Tg mutants.

### Defining the Tg Proteostasis Interactome

To identify protein-protein interactions potentially responsible for the aberrant processing of CH-associated mutations, we implemented an affinity purification – mass spectrometry (AP-MS) method coupled with tandem-mass-tag (TMT) labeling to allow for multiplexed identification and quantification of interacting proteins (Fig. 2A) (Plate *et al*, 2019). Transient interactions between Tg and PN components were captured using the cell-permeable cross-linker dithiobis(succinimidyl propionate) (DSP). We first sought to define the interactomes for WT Tg and each of the mutant variants (G2341R, L2284P, A2234D, and C1264R). We employed a mock AP using transfection of an untagged WT Tg control construct, or fluorescent proteins (EGFP or tdTomato) to delineate high confidence interactors from background proteins (Fig. 2A). We further optimized normalization methods and cutoffs to confidently identify interactors pertaining to protein folding, trafficking, and secretion likely to play a role in Tg processing (Fig. S2AB and Table S1) (Keilhauer *et al*, 2015; Chen *et al*, 2013). We identified interacting proteins for each individual Tg construct (Fig. S2C) and defined a cumulative list of 188 confidently identified interactors across all Tg constructs (Tables S2-3), we defined this list as the Tg interactome and focused on these proteins for subsequent analyses (Fig. 2B and Table S4). Using gene ontology (GO) enrichment analysis, 67% of the Tg interactome was enriched in components belonging to PQC within the secretory pathway (Fig 2C and Tables S3 and 5), including Hsp70/40 and Hsp90 chaperones and co-chaperones, N-glycosylation machinery, components involved in disulfide bond formation, ER-associated degradation (ERAD), as well as lysosomal and Golgi localized proteins (Chen *et al*, 2013).

**Figure 2.**
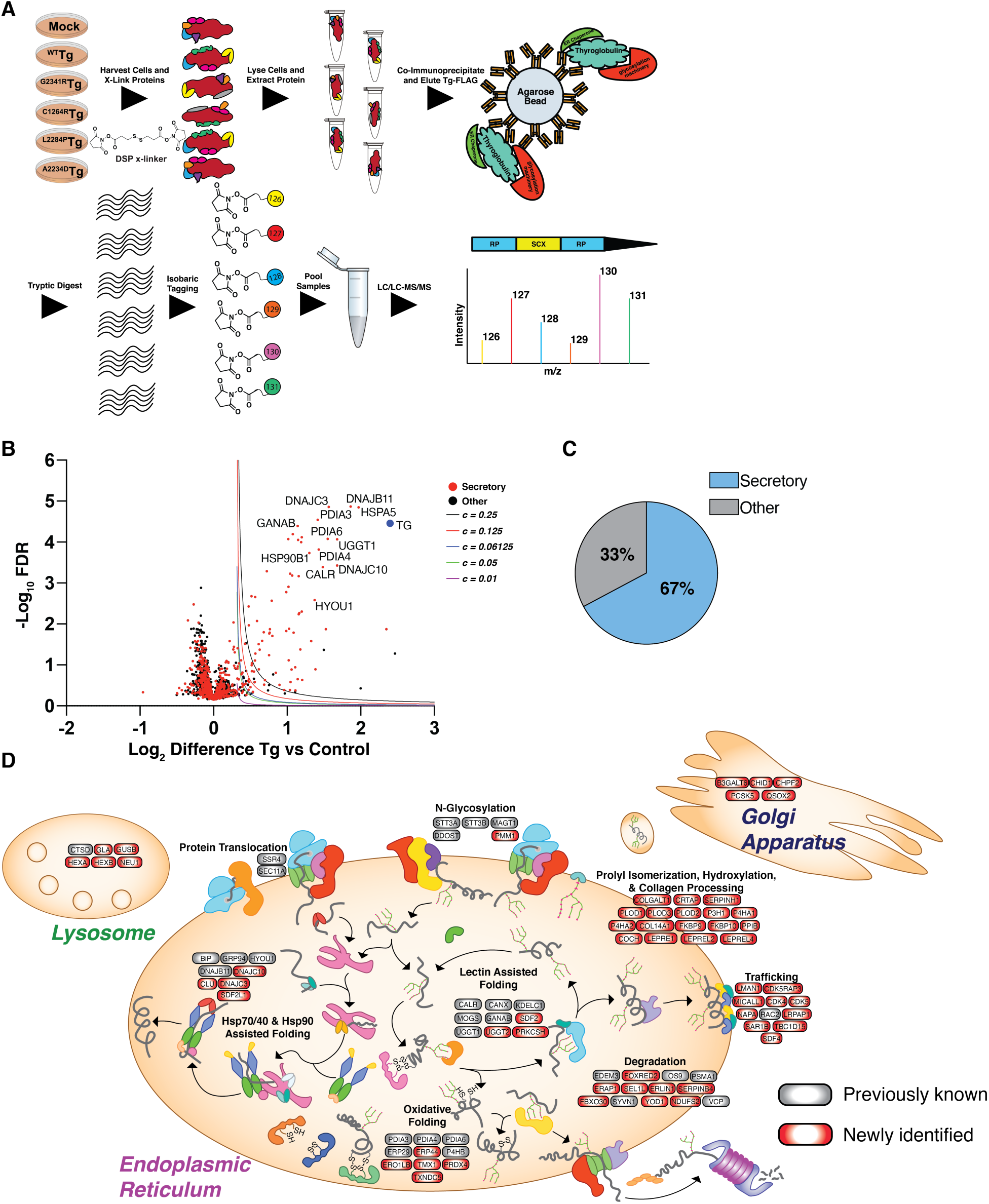
Defining the Tg interactome using multiplexed quantitative AP-MS. **A**. Schematic detailing the multiplexed quantitative interactomics workflow utilizing insitu crosslinking, affinity purification mass spectrometry (AP-MS), and tandem mass tags (TMT) for relative quantification of identified interactors to delineate interaction changes from WT to mutant variants. **B**. Volcano plot showing TMT enrichment ratios (log_2_ difference all Tg channels versus all mock channels) versus -log10 false discovery rate (Storey) for coimmunoprecipitated proteins of all Tg channels compared to all mock channels (n = 13 biological replicates). Variable cutoffs were used to optimize confident interactors of Tg. Optimization described in Fig. S2 and Table S1. Source data found in Table S4. **C**. Proteins found to be confident interactors with Tg are enriched within the secretory pathway. Source data found in Table S3. **D**. Schematic detailing newly identified Tg interactors (red) compared to previously publishes interactors (grey). Tg interactors are organized by biological function and organellar localization.

The Tg interactome identified here greatly expands the limited list of previously identified interactors. Importantly, our dataset also confirms previously known binding partners, such as GRP94, BiP, PDIA3, CANX, and CALR (Kim & Arvan, 1995; Menon *et al*, 2007). We were able to identify additional ER Hsp40 co-chaperones, including DNAJC3, DNAJB11, and DNAJC10, that can bind Tg directly and coordinate with the ER Hsp70 chaperone BiP to influence quality control decisions (Pobre *et al*, 2019; Behnke *et al*, 2016). Our data set is also rich in PN components involved in disulfide processing and formation. This enrichment is consistent with a strong dependence on oxidative folding pathways with Tg containing 122 cysteine residues and 61 disulfide bonds (Coscia *et al*, 2020). Only PDIA1/PDI, PDIA3/ERp57, PDIA4/ERp72, PDIA6/ERp5 and PDIA9/ERp29 were previously known to associate with Tg (Di Jeso *et al*, 2014; Baryshev *et al*, 2004; Menon *et al*, 2007), but our dataset additionally identified PDIA10/ERp44, TXNDC5/ERp46, and TMX1, among others (Figure 2B). ERAD associated factors, OS-9, EDEM3, SEL1L, which have been presumed to interact with Tg but not confirmed (Di Jeso & Arvan, 2016), along with new factors such as FOXRED2, and lysosomal components were also identified. The identification of lysosomal components suggest that autophagy may play a role in the degradation of some Tg constructs. Additionally, we detected previously known and novel interactors involved in glycoprotein folding and processing such as GANAB, LMAN1, UGGT1, as well as other lectins and glycan modifying enzymes. Overall, our analysis validated 28 previously identified Tg interactors and described 160 new PN interactions (Fig. 2D and Table S3).

### The Secretion Defect of Tg Mutants is Associated with Common Increases in Proteostasis Interactions

Next, we quantified interaction fold changes for the specific mutants relative to WT Tg to determine what factors may govern the aberrant PQC processing and secretion defects (Fig. 3A-B, Table S6, and Fig. S3A). Remarkably, when comparing CH-associated mutant interactomes with that of WT, many of the quantified interaction changes were similar across all CH-associated mutants. This held true particularly for factors involved in Hsp70/40 or Hsp90 assisted protein folding, including the Hsp70 and Hsp90 chaperones BiP and GRP94 along with co-chaperones DNAJB11 and DNAJC10. We observed similar increases with disulfide/redox-processing enzymes such as protein disulfide isomerases PDIA3, PDIA4, and PDIA6. We validated increased interactions between mutant Tg variants and GRP94, BiP, PDIA4, PDIA6, DNAJC10 by Co-AP followed by quantitative Western blot (Fig. S3B-C).

**Figure 3.**
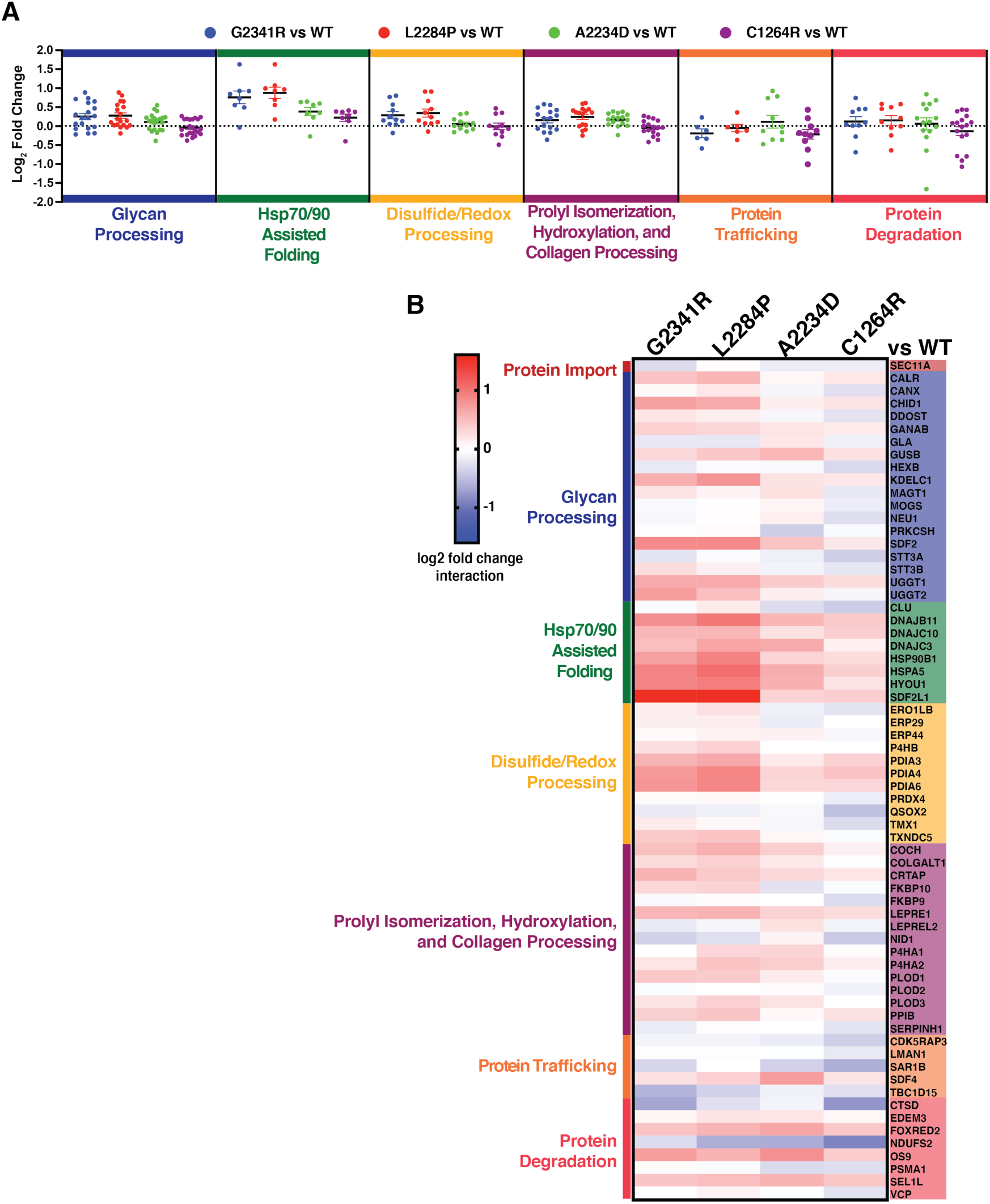
The secretion defect of Tg mutants is associated with both common and mutant specific changes in proteostasis interactions. **A**. Dot plots displaying aggregate interactome changes of proteostasis pathways between the different mutant Tg variants (G2341R, L2284P, A2234D, C1264R) compared to WT. Proteostasis factors are grouped based on biological function as in Fig. 3B and dots represent interaction changes for individual high-confidence interactors of Tg belonging to each group. Source data found in Table S5 and 6. **B**. Heatmap displaying altered interactions of mutant Tg variants with individual proteostasis components that were identified as high-confidence interactors. Interactors are grouped by biological function as in Fig. 3A. Source data found in Table S5 and 6.

Additionally, the enzymes responsible for marking and trafficking ER clients for ERAD including EDEM3, FOXRED2, OS9, and SEL1L all showed consistently increased interactions with Tg mutants compared to WT (Fig. 4A-D) (Christianson *et al*, 2008; Tang *et al*, 2014; Hirao *et al*, 2006; Bernasconi *et al*, 2008). This observation prompted us to test whether the CH-associated mutant Tg variants are instead degraded at a higher rate. To monitor potential changes in degradation rates of the Tg constructs we employed a cycloheximide (CHX) chase assay (Fig. S4A). Approximately one third of WT Tg was secreted after 4 hours (Fig. 4E and Fig. S4B). As previously noted, none of the CH-associated Tg mutants were secreted. When monitoring Tg degradation, all constructs showed similar rates of degradation on the scale of 30 - 40% after 4 hours of CHX treatment (Fig. 4E and Fig. S4B). This degradation rate is consistent with prior studies (Tokunaga *et al*, 2000) and indicated that despite increased targeting of mutant Tg towards ERAD, the degradation rates are unaffected. To investigate why degradation rates are unchanged for the mutant variants, we next looked at downstream proteostasis factors involved in ERAD after retrotranslocation of proteins into the cytosol. We detected Tg interactions with VCP/p97, the ATPase involved in extracting substrates from the ER, as well as several subunits of the proteasome (Fig. 4A,F). Interactions between Tg mutant variants and VCP are mostly unchanged relative to WT, and proteasome subunits were consistently decreased for all mutants. Together, these results suggest that, while CH-associated Tg mutants are recognized and processed for ERAD, the proteins may not be properly retrotranslocated for subsequent proteasomal degradation.

**Figure 4.**
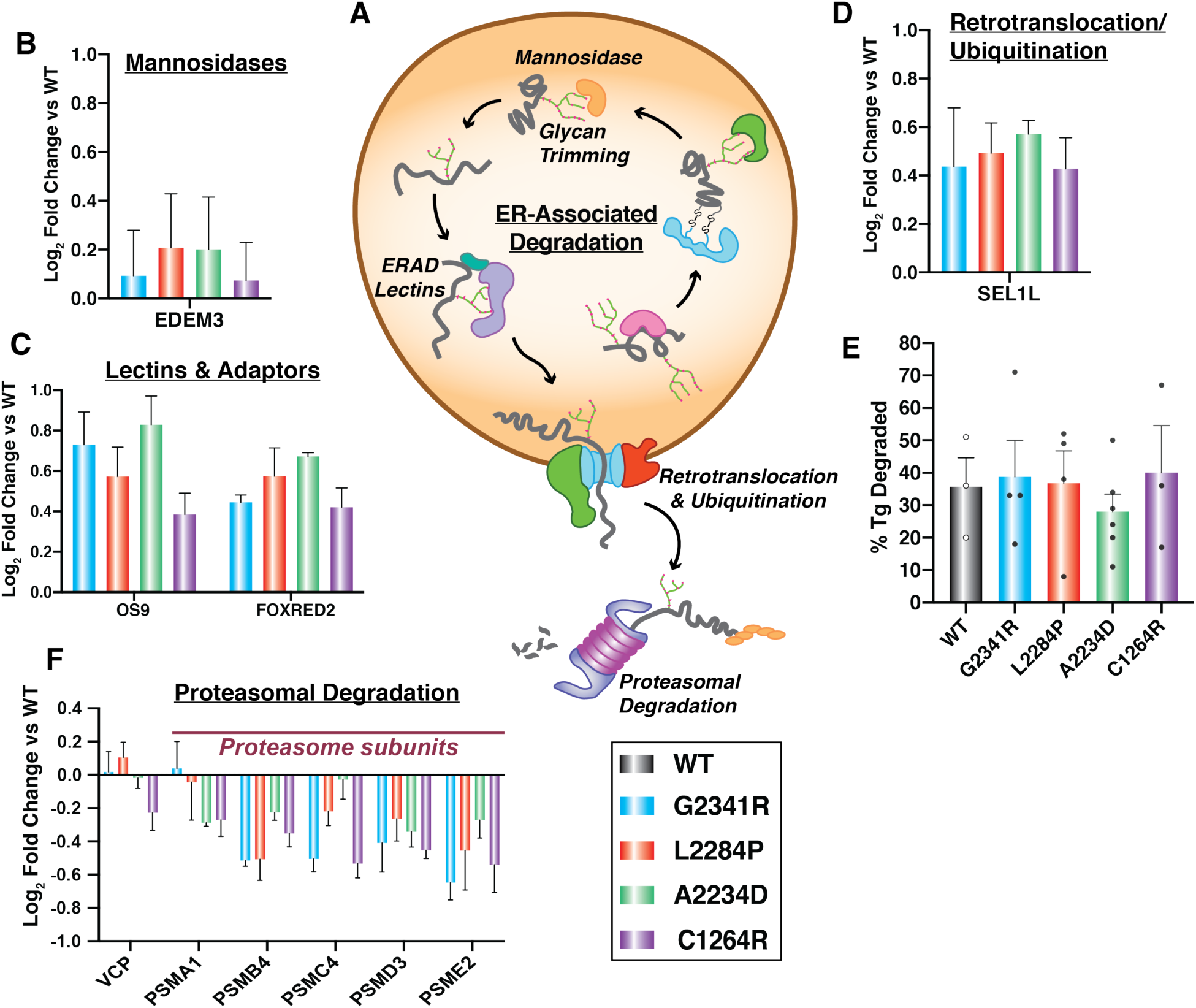
Tg mutants are increasingly routed towards ER-associated degradation machineries but not degraded at faster rates. **A**. Schematic detailing the ERAD pathway: degradation factors targeting proteins through glycan trimming, subsequent retrotranslocation and ubiquitination, followed by proteasome-mediated degradation in the cytosol. **B – D**. Interaction changes of Tg mutants compared to WT with individual ERAD factors that were identified as high-confidence interactors of Tg. Error bars show SEM. **B**. Mannosidases responsible for glycan trimming. **C**. An ERAD specific lectin (OS-9) and another ERAD factor (FOXRED2) **D**. A subunit of the retrotranslocation complex. **E**. Plot showing the percentage of Tg degradation measured in HEK293^DAX^ cells 4 hours after treatment with 50µg/mL of cycloheximide to block new protein translation. Error bars show SEM for 3-6 biological replicates. There is no significant difference in degradation for any of the Tg mutants compared to WT. Student’s parametric T test was used to determine significant changes (p < 0.05). G2341R: p = 0.849, L2284P: p = 0.942, A2234D: p = 0.464, C1264R: p = 0.813. Representative Western blots for the CHX chase experiments are shown in Fig. S4B. **F**. Interaction changes of Tg mutants compared to WT for cytosolic proteins involved in proteasome-mediated protein degradation. Error bars shows SEM.

Overall, the interactomics data suggest that CH-associated mutant Tg display common PQC defects linked to prolonged chaperoning facilitated largely by Hsp70/40, Hsp90, and disulfide/redox folding pathways, as well as increased associations with ER luminal ERAD components. Interestingly, in many cases interaction fold changes were slightly higher for all ChEL domain Tg mutants, G2341R, L2284P, and A2234D than for the C1264R mutant occurring in the hinge/flap region. This parallels our and previous secretion data exhibiting residual C1264R-Tg secretion (Lee *et al*, 2011) and suggest that the Tg PQC defects are more profound when mutations occur in the ChEL domain (Fig. 3B, Table S6, and Fig. S1A-C).

### Tg Mutants Show Distinct Changes in Engagement with N-linked Glycosylation Pathways

While common changes in PN interactions across CH-associated mutants provide new insights on conserved Tg processing mechanisms, we wanted to further explore the data to investigate whether any mutation-specific PN interaction changes occurred that may define unique PQC defects associated with each mutation. We detected subtle, yet striking deviations across CH-associated Tg interactions with PN components involved in N-glycosylation and lectin folding (Fig. 5A-E). The membrane bound lectin calnexin (CANX) showed modestly increased interactions with G2341R and L2284P variants, yet decreased interactions in the case of C1264R and A2234D. Intriguingly, interactions with calreticulin (CALR), the soluble paralog of CANX, were also more strongly increased for the G2341R and L2284P variants, along with PDIA3, which has been shown to specifically bind both CANX or CALR in complex (Kozlov *et al*, 2006; Lamriben *et al*, 2016) (Fig. 5B). The glucosyltransferases UGGT1 and UGGT2 showed similarly divergent interaction changes (Fig. 5C). In the case of UGGT2, all mutations in the ChEL domain exhibited increased interactions while C1264R showed modestly decreased interactions. Yet, UGGT1 interactions across all mutations were increased. In line with this observation was the increased interactions across all mutations with GANAB, the catalytic subunit of the heterodimeric ER glucosidase II complex responsible for sequentially cleaving the two innermost glucose residues of the ER associated N-linked high-mannose oligosaccharide precursor – a necessary process required for client proteins to properly enter the CANX/CALR lectin folding pathway (Fig. 5D) (Martiniuk *et al*, 1985; Lamriben *et al*, 2016). While these findings revealed some unique mutation specific changes, all of these interactions occur downstream of N-glycosylation by the oligosaccharyltransferase (OST) complex, prompting us to compare potential interaction changes with the OST complex (Braunger *et al*, 2018). We noticed mutation specific changes in engagement with the two different OST isoforms. In one isoform containing the catalytic STT3A subunit, the OST is largely associated with the translocon channel and facilitates co-translational glycosylation of ER client proteins (Ruiz-Canada *et al*, 2009). Most CH-associated Tg mutants showed modestly decreased interactions with the STT3A catalytic subunit relative to WT (Fig. 5E). The other OST isoform containing STT3B is largely associated with post-translational glycosylation of ER client proteins (Ruiz-Canada *et al*, 2009). Here, G2341R and L2284P exhibited modestly increased or unchanged interactions with STT3B while A2234D and C1264R showed decreased interaction (Fig. 5E). We confirmed by Co-AP and Western blot that A2234D and C1264R displayed decreased interactions with STT3B compared to WT and that these changes were distinct from G2341R and L2284P (Fig. S4C).

**Figure 5.**
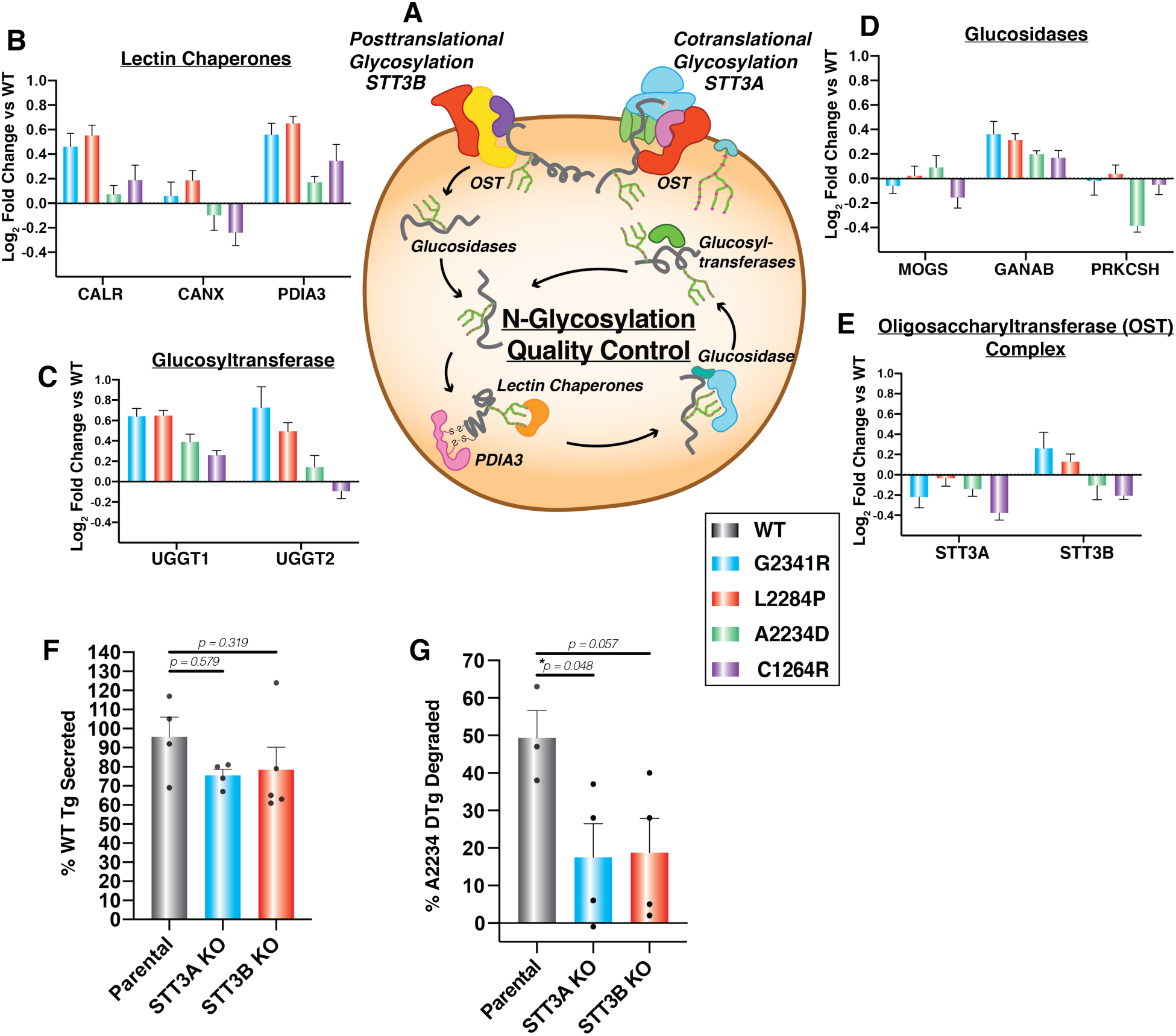
Perturbation of N-linked glycosylation distinctly impacts A2234D Tg. **A**. Schematic detailing the N-glycosylation and lectin-mediated folding pathway. Glycosylation is carried out by two distinct OST complexes containing STT3A or STT3B as catalytic subunits. Glucosidases then trim terminal glucose residues, lectin chaperones (CANX, CALR) and PDIA3 assist in folding, followed by further glucose trimming. Subsequently, glucosyltransferases (UGGT1, UGGT2) serve as quality control sensors to re-glucosylate improperly folded proteins for iterative chaperoning cycles. **B – E**. Interaction changes of Tg mutants compared to WT with individual N-glycosylation quality control factors that were identified as high-confidence interactors of Tg. Error bars show SEM. **B**. Lectin chaperones and lectin-associated protein disulfide isomerase PDIA3. **C**. Glucosyltransferases sensing misfolded proteins. **D**. Glucosidases involved in glycan trimming. **E**. Catalytic subunits of the OST complex STT3A responsible for cotranslational glycosylation and STT3B responsible for posttranslational glycosylation. **F**. Comparison of WT Tg secretion in parental, STT3A, or STT3B KO HEK293T cells. WT Tg was transiently transfected into the respective cells and newly synthesized proteins were metabolically labeled for 30 min with ^35^S and then chased with unlabeled media. ^35^S-labeled protein was quantified in the media and lysate after 4 hours. % Tg secreted was calculated as Tg_media, 4h_ /(Tg_lysate, 0h_+Tg_media, 0h_.) Error bars represent SEM of 3-4 biological replicates. STT3A or STT3B KO do not significantly alter WT secretion. Student’s parametric t test was used to determine significant (p < 0.05) changes in Tg secretion and p values are indicated. Representative autoradiograms are shown in Fig. S4J. **G**. Comparison of A2234D Tg degradation in parental, STT3A, or STT3B KO HEK293T cells. A2234D Tg was transiently transfected into the respective cells and subjected to the same ^35^S-pulse labeling scheme as in F % Tg degraded was calculated as 1(Tg_lysate, 4h_ /Tg_lysate, 0h_). Error bars represent SEM of 3-4 biological replicates. Representative autoradiograms are shown in Fig. S4I. Student’s parametric t test was used to determine significant (p < 0.05) changes in Tg degradation and p values are indicated.

**Figure 6.**
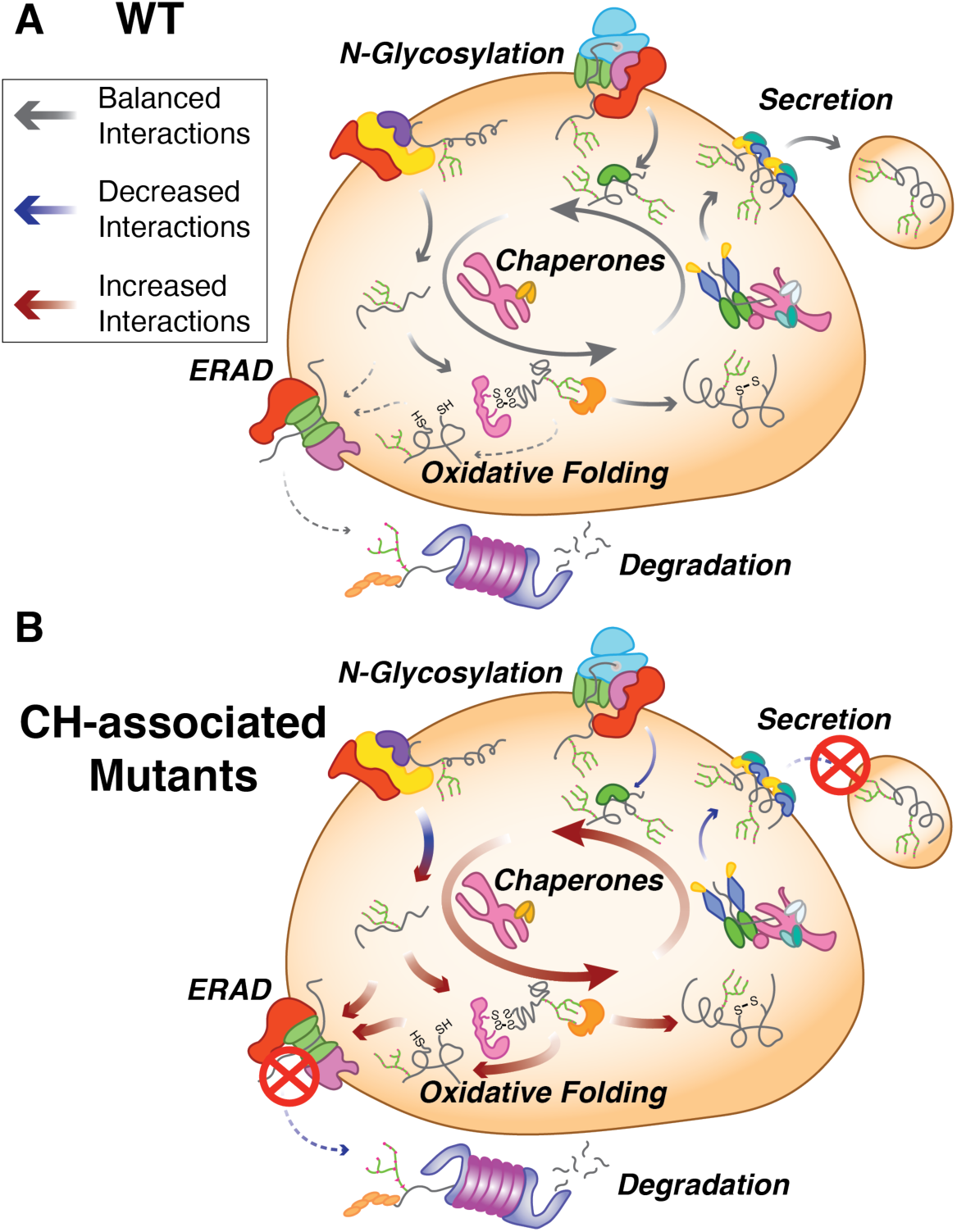
Model for common and mutant-specific proteostasis interactome changes mediating the secretion defect of CH-associated Tg variants. **A**. In the case of WT Tg processing (top) the delicate balance of proper chaperoning, post-translational modification and secretion are maintained to provide sufficient Tg processing, secretion, and subsequent hormone production (indicated by the arrow size and color code denoted in the key). **B**. In the case of secretion-defective, CH-associated Tg mutants (bottom) this balance between chaperoning is shifted in such a way that increased chaperoning and engagement with oxidative folding enzymes, possibly stemming from altered engagement with the OST complex dominates Tg processing (indicated by the arrow size and color code denoted in the key). Additionally, mutant Tg is increasing marked for degradation, yet inefficient retrotranslocation or decreased engagement by the proteasome leads to degradation rates remaining consistent compared to WT.

Overall, our findings reveal distinct PQC defects for the different CH-associated Tg mutants and their engagement with the OST complex and downstream processing through the CANX/CALR lectin folding pathway (Lamriben *et al*, 2016).

### Perturbation of N-linked Glycosylation Distinctly Impacts Tg Mutants

The OST complex is most upstream in the N-glycosylation and lectin folding pathway (Fig 5A). Therefore, we chose to focus on elucidating the role of the two different STT3A and STT3B dependent OST isoforms on downstream Tg processing. To assess the functional implications of these changes with OST engagement identified in our dataset, we monitored the effects of isoform-specific knockouts of STT3A or STT3B Nglycosylation machinery on Tg secretion and degradation (Kelleher *et al*, 2003; Ruiz-Canada *et al*, 2009; Cherepanova & Gilmore, 2016). We transfected Tg variants into HEK293T STT3A^−/−^ and STT3B^−/−^ knockout cell lines and followed Tg processing via cycloheximide (CHX) chase assay and ^35^S pulse-chase labeling (Fig. S4A,D). Knockout of either OST subunit did not abolish WT Tg secretion nor rescue secretion of any CH mutants (Fig. 5F, S4E). On the other hand, small decreases in secretion for WT Tg occurred in the STT3A and STT3B KO cells when measured via CHX assay (Fig. S4F-G). We then went on to quantify degradation rates for WT Tg and the CH-associated mutants (Fig. S5F,H). For all Tg constructs, degradation rates were not significantly changed in the STT3A or STT3B KO cells compared to the parental cells, yet we noticed a small, but variable, increase in degradation for G2341R in the presence of either STT3A or STT3B KO and a small, but insignificant, attenuation of degradation of A2234D in STT3A knockout cells. Given the high variability in degradation measurements using the CHX chase assay, we employed the ^35^S pulse-chase labeling scheme to mitigate any complications resulting from inhibiting protein synthesis. Using the ^35^S assay, STT3A and STT3B KO cells resulted in significantly reduced degradation rates for A2234D Tg, from 50% to 20% degraded after 4 hours (Fig 5G, Fig. S4I). Furthermore, STT3A and STT3B KOs did not significantly alter WT Tg degradation (Fig. S4JK). These results suggest that in the case of A2234D, engagement with both OST isoforms is necessary for proper entry into the CANX/CALR lectin cycle and degradation initiated by glycoprotein quality control within the ER.

## DISCUSSION

Most CH-associated Tg missense mutant variants present with very similar phenotypes resulting in inefficient trafficking and loss of secretion. The majority of these CH mutant variants have been reported to be retained within the ER, interact with canonical PN components, and induce the unfolded protein response (UPR), which acts to adjust ER quality control capacity in response to ER stress (Baryshev *et al*, 2004; Plate & Wiseman, 2017). However, there has been little investigation into 1) identifying molecular mechanisms involved in the inefficient folding and trafficking of these mutant proteins, and 2) developing or exploring therapeutic avenues aimed at rescuing the secretion and subsequent hormone production from these mutants (Kanou *et al*, 2007; Caputo *et al*, 2007; Pardo *et al*, 2009; Hishinuma, 1999; Baryshev *et al*, 2004; Menon *et al*, 2007; Kim *et al*, 1996). Here we utilized a multiplexed quantitative interactomics platform to describe the PN dependencies of several CH-associated Tg variants with mutations clustering in two different domains of Tg. Our analysis of Tg interactomes reveals common PQC defects that are involved in the loss of Tg secretion, but we also identify unique dependencies that may suggest PQC mechanisms that are specific to individual mutations.

Using the quantitative interactomics profiling method, we were able to identify previously documented Tg interactors such as BiP, GRP94, CANX, CALR, and PDIA3, but also greatly expand our knowledge of additional cochaperones, lectins, trafficking and degradation factors that influence PQC activity and subsequent Tg processing. Previous work has provided insights on the implications of some of these interactions as BiP overexpression decreases WT Tg secretion (Muresan & Arvan, 1998). The iterative binding cycles between Tg with BiP and co-chaperones likely results in overall increased retention of Tg and therefore blocks partitioning of Tg to necessary trafficking components (Awad *et al*, 2008). Additionally, our analysis identified a number of Hsp40 co-chaperones, which can direct chaperone pathways to assume pro-folding or pro-degradation roles (Behnke *et al*, 2016; Oikonomou & Hendershot, 2020). PDIA4 has also been identified as a key interactor as it has been shown to bind mutant Tg and form coaggregates retained within the ER (Menon *et al*, 2007). We identified increased interaction between PDIA4 and all secretion-deficient Tg mutants in this study, but we also identified stronger engagement with many other protein disulfide isomerase protein family members and factors involved in oxidative protein folding. Several PDI family members were already previously shown to form transient mixed-disulfide-linked intermediates with Tg highlighting their involvement in Tg folding (Di Jeso *et al*, 2014; Baryshev *et al*, 2004; Menon *et al*, 2007). The increased interactions with these PDIs and additional factors involved in disulfide bond formation may be responsible for intracellular retention and co-aggregation with destabilized Tg variants, which may be mediated through non-resolved mixed-disulfide bond intermediates (Kim *et al*, 1993; Menon *et al*, 2007).

Our results show that the CH-associated Tg mutants not only engage many of the chaperoning and oxidative protein folding pathways to a greater extent, but also suggest altered interactions with ERAD and N-glycosylation components that may be responsible for the retention or aggregation within the ER. Prior work has shown that ERAD of Tg is suppressed upon the inhibition of mannosidase I (MAN1B1) (Tokunaga *et al*, 2000). MAN1B1 is known to trim the outermost alpha-1,2-linked mannose residue of the high-mannose ER-associated glycan, followed by subsequent trimming of inner mannose residues, a key step within the glycoprotein ERAD process (Avezov *et al*, 2008). While we did not identify MAN1B1 in our dataset, other mannosidases of the ER degradation enhancing alpha-mannosidase like (EDEM) family associated with ER stress and UPR activation were identified in the current study (Molinari *et al*, 2003; Oda *et al*, 2003). These factors exhibited predominantly increased interactions with CH-associated mutants relative to WT Tg, along with accessory proteins such as SEL1L, FOXRED2, OS-9, and other vital glycoprotein ERAD components (Christianson *et al*, 2008; Bernasconi *et al*, 2008; Christianson *et al*, 2012). This increased engagement with ERAD targeting factors seemed paradoxical considering that mutant Tg variants were not degraded to a greater extent than WT Tg. However, the mutant Tg variants may be recognized as destabilized and targeted for ERAD, but not efficiently retrotranslocated or recognized by the proteasome to initiate degradation (Nakatsukasa *et al*, 2013). In support of this model, we observed decreased interactions between Tg mutant variants and several subunits of the proteasome. Instead, any of the above-mentioned chaperone/PDI-mediated retention or co-aggregation mechanisms could compete with retrotranslocation and degradation. It has been shown that protein aggregation propensity and ERAD targeting are tightly linked (Sun & Brodsky, 2018). Ultimately, Tg must be degraded as indicated by prior work and from our CHX and pulse-chase experiments, likely through residual ERAD activity and also autophagy (Menon *et al*, 2007; Tokunaga *et al*, 2000). Consistent with this, we identified a number of lysosomal protein factors (Fig. 2D and Fig. S5). Consequently, further investigations into the degradation components and pathways facilitating Tg degradation would be of interest to examine, particularly cross talk and timing between ERAD and ER-phagy (Schuck *et al*, 2014; Pohl & Dikic, 2019; Carvalho *et al*, 2006; Oikonomou & Hendershot, 2020; Sun & Brodsky, 2019).

The identification of altered interactions between CH-associated Tg variants and the components of the OST complex involved in protein N-glycosylation provides new insights into the distinct misprocessing defects for individual CH-associated Tg mutants. While the disruptions in OST complex interactions do not completely explain why these Tg mutants are unable to exit the ER, these changes occur for the most upstream enzymes mediating Tg post-translational processing suggesting an important effect. Importantly, subtle changes in glycosylation patterns could have profound influences on binding of lectin-chaperones, glycan processing enzymes, or lectin-associated oxidoreductases (Hammond *et al*, 1994; Kozlov *et al*, 2006; Martiniuk *et al*, 1985; Lamriben *et al*, 2016).

The changes in PN engagement could be a consequence of changes in Tg secondary and tertiary structure of folding intermediates that then lead to altered engagement with the OST complex. Changes in secondary structure have been loosely predicted for some Tg mutants including extension or reduced stretch of α -helix or β -sheet structure, along with the formation of a β -sheet (Pardo *et al*, 2008). Interestingly, the recent Cryo-EM structure of Tg showed that the three mutations, A2234D, L2284P, and G2341R cluster into a small region of the ChEL domain, suggesting that all three mutations could destabilize the structure at a similar location (Fig. S1B). Despite this co-localization, the ChEL mutants displayed several distinct interactome changes and only A2234D could be rescued from degradation by knockout of individual OST isoforms. The same cryo-EM structure also revealed that a glycosylation site at N2013 may play a key role in stabilizing the Tg dimer (Coscia *et al*, 2020). Investigations into which OST complex is responsible for N2013 glycosylation and how these ChEL domain mutations may change the glycan occupancy would be of particular interest. On the other hand, the C1264R mutation is localized distantly from the ChEL domain in the hinge/flap region, presumably disrupting a local disulfide bond with C1245 and resulting in structurally distinct folding defects (Fig. S1D). Nonetheless, interaction changes with the PN for C1264R are largely similar to the ChEL domain mutants, albeit to a lower magnitude, highlighting that mutations in distinct regions of the protein can produce common PQC deficiency that result in loss of protein secretion. Additionally, disulfide bond formation and N-glycosylation are competing reactions within the ER, further complicating Tg processing (Allen *et al*, 1995).

While PQC pathways are unable to facilitate complete folding and secretion of the Tg mutants, folding and processing is clearly attempted prior to timely degradation taking place. It may be the case for A2234D Tg that in the absence of either STT3A or STT3B in the KO cells, proper entry into the CANX/CALR cycle is disrupted, ultimately leading to retention within the ER and decreased degradation. The decreased degradation rate may stem from an inability of the PN to recognize A2234D Tg for glycoprotein ERAD due to aberrant glycosylation, or aberrant glycosylation may lead to the preferential aggregation of A2234D Tg, allowing it to escape ERAD (Lamriben *et al*, 2016; Oikonomou & Hendershot, 2020). Paradoxically, the other three mutant Tg variants studied here do not exhibit as stark a dependency on the presence of both OST isoforms for efficient degradation, although these mutants also displayed altered engagement with OST components in the interactomics dataset and two of the studied mutations fall into the same structural region of the ChEL domain as A2234D. Overall, our results highlight that altered proteostasis interactions with Tg variants can have subtle, yet significant functional outcomes that are highly specific to localization and nature of the destabilizing mutation. The cryo-EM structure of human Tg and future structural studies on mutant Tg variants could enable further insights into what structural changes influence the engagement of proteostasis factors (Coscia *et al*, 2020).

The modulation of PN components or entire pathways has shown recent promise as a therapeutic strategy to combat a number of protein folding diseases (Plate & Wiseman, 2017; Hetz *et al*, 2019; Ryno *et al*, 2014; Plate *et al*, 2016; Chen *et al*, 2014). By using quantitative multiplexed interactome proteomics we identified specific PN components that may act as therapeutic targets for rescuing Tg secretion. A similar method has been used to investigate the molecular basis of activating transcription factor 6 (ATF6) dependent regulation of immunoglobulin light chain secretion in (AL) amyloidosis (Plate *et al*, 2019). Methods to pharmacologically target the UPR may be further applicable to rescuing Tg secretion. Many of the CH-associated mutations presented here naturally activate the UPR (Caputo *et al*, 2007; Baryshev *et al*, 2004; Kim *et al*, 1996). Therefore, pharmacologic modulation of UPR activity to regulate the abundance of ER proteostasis factors in a coordinated manner may potentially act to restore mutant Tg secretion. In the case of amyloidogenic light chain proteins, overexpression of UPR-regulated chaperones, in particular BiP and GRP94, was able reduce the secretion of an aggregation-prone protein variant (Cooley *et al*, 2014; Plate *et al*, 2019). Similar effects were also observed for a model aggregation-prone polyQ protein as cytosolic heat shock activation attenuated intracellular aggregation, and cellular toxicity (Ryno *et al*, 2014). In contrast, the increased surveillance of destabilized Tg variants by chaperoning and oxidative folding pathways are likely directly implicated in the loss of protein secretion. Here, UPR activation potentially further exacerbates the secretion defects for Tg mutants by increasing the abundance of relevant PN components and promoting increased intracellular interactions. Consequently, reducing the engagement between mutant Tg variants and the identified chaperoning, oxidative folding, and ERAD targeting pathways could be a viable strategy to restore mutant Tg secretion (Gallagher & Walter, 2016; Cross *et al*, 2012; Plate & Wiseman, 2017). Our quantitative proteostasis interactome map forms the framework for the identification of single PN components or entire pathways as viable drug targets geared towards rescuing Tg secretion. Future studies directed at disrupting the individual protein interactions or reducing PN capacity in a coordinated manner through pharmacologic inhibition of UPR signaling pathways could reveal the impact on rescue of CH-associated mutant Tg secretion.

## AUTHOR CONTRIBUTIONS

M.T.W. and L.P. designed experiments. M.T.W., L.K., and L.P. performed the experiments and analyzed data. M.T.W. and L.P. wrote the manuscript.

## DATA DEPOSITION

The mass spectrometry proteomics data have been deposited to the ProteomeXchange Consortium via the PRIDE partner repository with the dataset identifier PXD018379.

## ACKNOWLEDGEMENTS

We thank Dr. Renã Robinson (Vanderbilt University) for the use of mass spectrometry resources and Dr. Reid Gilmore (University of Massachusetts Medical Center) for providing STT3A and STT3B knockout cell lines. This work was funded by the Vanderbilt Institute of Chemical Biology Fellowship, Vanderbilt Chemistry-Biology Interface Training Program (NIGMS, 5T32GM065086), National Science Foundation Graduate Research Fellowship Program (M.T.W.), and an NIGMS R35 award (1R35GM133552).

## METHODS

### Plasmids and Antibodies

FLAG-tagged (FT)-Tg in pcDNA3.1+/C-(K)-DYK plasmid was purchased from Genscript (Clone ID OHu20241). Site-directed mutagenesis was then performed to engineer FTG2341R, FT-L2284P, FT-C1264R, FT-A2234D, and untagged Tg plasmids (Table S7). Primary antibodies were acquired from commercial sources and used at the indicated dilutions in immunoblotting buffer (5% bovine serum albumin (BSA) in Tris-buffered saline pH 7.5, 0.1% Tween-20, and 0.1% sodium azide). Mouse monoclonal antibodies were used for the detection of KDEL (1:1000, Enzo Life Sciences, ADI-SPA-827), M2 anti-FLAG (1:1000, Sigma Aldrich, F1804). Polyclonal rabbit antibodies were used to detect Calnexin (1:1000, GeneTex, GTX109669), PDIA4 (1:1000, Proteintech, 14712-1AP) DNAJC10 (1:500, Proteintech, 13101-1-AP), thyroglobulin (1:1000, Proteintech, 21714-1-AP), UGGT1 (1:1000, Proteintech, 14170-1-AP). STT3A (1:2000, Proteintech, 12034-1-AP) and STT3B (1:2000, Proteintech, 15323-1-AP) Secondary antibodies were obtained from commercial sources and used at the indicated dilutions in 5% milk in Trisbuffered saline pH 7.5, 0.1% Tween-20 (TBS-T): Goat anti-mouse Starbright700 (1:10000, Bio-Rad,12004158), Goat anti-rabbit IRDye800 (1:10000, LI-COR, 926-32211), Goat anti-rabbit Starbright520 (1:10000, Bio-Rad,12005869).

### Cell Culture and Transfections

HEK293^DAX^ cells (Shoulders *et al*, 2013), HEK293T, and STT3A or STT3A KO HEK293T cells (Cherepanova & Gilmore, 2016) were grown in Dulbecco’s modified Eagle’s medium (DMEM) supplemented with 10 % fetal bovine serum (FBS), 1% L-glutamine (200mM), 1% penicillin (10,000U) / streptomycin (10,000 µg/ml). Generally, cells were transiently transfected with respective Tg expression plasmids using a calcium phosphate method.

### Affinity Purification and MS Sample Preparation

A fully confluent 10cm tissue culture plate (approximately 10^7^ cells) was used per condition. Cells were harvested by washing with PBS and incubating with 1mM EDTA in PBS on ice. A cell scraper was then used to dislodge cells. Cells were harvested, washed once with PBS, and treated with 0.5mM dithiobis(succinimidyl propionate) (DSP) (Thermo Scientific, PG82081) in PBS for 30 minutes at room temperature while rotating. Crosslinking was quenched by addition of 100mM Tris pH 7.5 for 15 minutes. Lysates were prepared by lysing in RIPA buffer (50mM Tris pH 7.5, 150mM NaCl, 0.1% SDS, 1% Triton X-100, 0.5% deoxycholate and protease inhibitor cocktail (Roche, 4693159001)) and protein concentration was normalized. Cell lysates were then precleared on 4B sepharose beads (Sigma, 4B200) at 4°C for 1 hour while rocking. Precleared lysates were immunoprecipitated with M2 anti-flag agarose resin (Sigma, A2220) or G1 Anti-DYKDDDDK affinity resin (GenScript, L00432) overnight at 4°C while rocking. Resin was washed four times with RIPA buffer. Proteins were eluted twice in 75uL elution buffer (2% SDS, 1mM EDTA, in PBS) by heating at 95°C for 5 minutes. Eluted samples were precipitated in methanol/chloroform, washed three times with methanol, and air dried. Protein pellets were then resuspended in 3uL 1% Rapigest SF Surfactant (Waters, 186002122) followed by the addition of 10uL of 50mM HEPES pH 8.0, and 32.5uL of H_2_O. Samples were reduced with 5mM tris(2-carboxyethyl)phosphine (TCEP) (Sigma, 75259) at room temperature for 30 minutes and alkylated with 10mM iodoacetimide (Sigma, I6125) in the dark at room temperature for 30 minutes. 0.5 ug of Trypsin (Promega, V511A or Thermo Scientific, PI90057) was then added and incubated for 16-18 hours at 37°C, shaking at 700rpm. Peptides were reacted with TMT sixplex reagents (Thermo Fisher, 90066) in 40% v/v acetonitrile and incubated for one hour at room temperature. Reactions were quenched by the addition of ammonium bicarbonate (0.4% w/v final concentration) and incubated for one hour at room temperature. TMT labeled samples for a given experiment were then pooled and acidified with 5% formic acid (Fisher, A117, v/v). Samples were concentrated using a speedvac and resuspended in buffer A (95% water, 4.9% acetonitrile, and 0.1% formic acid, v/v/v). Cleaved Rapigest SF surfactant was removed by centrifugation for 30 minutes at 21,100 x g.

### Mass Spectrometry and Interactome Characterization

MudPIT microcolumns were prepared as previously described (Fonslow *et al*, 2012). Peptide samples were directly loaded onto the columns using a high-pressure chamber. Samples were then washed for 30 minutes with buffer A (95% water, 4.9% acetonitrile, 0.1% formic acid v/v/v). LC-MS/MS analysis was performed using a Q-Exactive HF (Thermo Fisher) or Exploris480 (Thermo Fisher) mass spectrometer equipped with an Ultimate3000 RSLCnano system (Thermo Fisher). MudPIT experiments were performed with 10uL sequential injections of 0, 10, 30, 60, and 100% buffer C (500mM ammonium acetate in buffer A), followed by a final injection of 90% buffer C with 10% buffer B (99.9% acetonitrile, 0.1% formic acid v/v) and each step followed by a 130 minute gradient from 5% to 80% B with a flow rate of either 300 or 500nL/minute on a 20cm fused silica microcapillary column (ID 100 um) ending with a laser-pulled tip filled with Aqua C18, 3um, 100 Å resin (Phenomenex). Electrospray ionization (ESI) was performed directly from the analytical column by applying a voltage of 2.0 or 2.2kV with an inlet capillary temperature of 275°C. Using the Q-Exactive HF, data-dependent acquisition of mass spectra was carried out by performing a full scan from 300-1800 m/z with a resolution of 60,000. The top 15 peaks for each full scan were fragmented by HCD using normalized collision energy of 35 or 38, 0.7 m/z isolation window, 120 ms maximum injection time, at a resolution of 15,000 scanned from 100 to 1800 m/z and dynamic exclusion set to 60s. Using the Exploris480, data-dependent acquisition of mass spectra was carried out by performing a full scan from 400-1600m/z at a resolution of 120,000. Top-speed data acquisition was used for acquiring MS/MS spectra using a cycle time of 3 seconds, with a normalized collision energy of 36, 0.4m/z isolation window, 120ms maximum injection time, at a resolution of 30000 with the first mass (m/z) starting at 110. Peptide identification and TMT-based protein quantification was carried out using Proteome Discoverer 2.3 or 2.4. MS/MS spectra were extracted from Thermo Xcalibur .raw file format and searched using SEQUEST against a Uniprot human proteome database (released 05/2014). The database was curated to remove redundant protein and splice-isoforms, and supplemented with common biological MS contaminants. Searches were carried out using a decoy database of reversed peptide sequences and the following parameters: 10ppm peptide precursor tolerance, 0.02 Da fragment mass tolerance, minimum peptide length of 6 amino acids, trypsin cleavage with a maximum of two missed cleavages, dynamic methionine modification of 15.995 Da (oxidation), static cysteine modification of 57.0215 Da (carbamidomethylation), and static N-terminal and lysine modifications of 229.1629 Da (TMT sixplex). SEQUEST search results were filtered using Percolator to minimize the peptide false discovery rate to 1% and a minimum of two peptides per protein identification. TMT reporter ion intensities were quantified using the Reporter Ion Quantification processing node in Proteome Discoverer 2.3 or 2.4 and summed for peptides belonging to the same protein.

### Interactome Characterization and Pathway Enrichment Analysis

To identify true interactors from non-specific background TMT intensities first underwent a log_2_ transformation, were then median normalized using the formula: 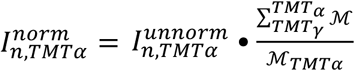. Here, 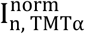 and 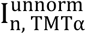 are the unnormalized and normalized TMT intensities for a given protein *n* found in TMT channels α - *γ*, and ℳ is the median TMT intensity value for TMT channels α - *γ*. TMT ratios were then calculated between respective Tg AP and control TMT channels using formula: 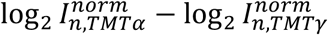. The mean of log_2_ interaction differences was then calculated across the multiple LC-MS batches (Fig. S2A). Significance of interaction differences was then calculated using a paired, parametric, two tailed t-test of 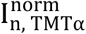, and multiple testing correction via FDR estimation (Storey & Tibshirani, 2003). A previously described method was used to delineate true interactors from non-specific background (Keilhauer *et al*, 2015). In short, the function *y* = *c*/(*x* – *x*0) was used to, where c = curvature and *x*0 = minimum fold change, set as one standard deviation of the of the Tg-containing TMT channel used for comparison. The *c* parameter was optimized to separate true interactors from false positives (Fig. S2C and Table S1). Tg interactors were identified for WT and mutant Tg individually. A cumulative list of identified interactors was then used for WT vs mutant Tg comparisons. To compare WT vs mutant Tg interactors TMT intensities were normal-ized using formula: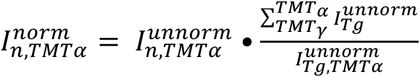. Here, 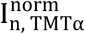 and 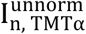 are the unnormalized and normalized TMT intensities for a given protein *n* found in TMT channels α -*γ*, and 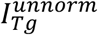 is the unnormalized TMT intensity value for Tg in a given TMT channel α -*γ*. For pathway enrichment analysis of identified protein, EnrichR was used and GO Cellular Component 2018 terms were used to differentiate secretory pathway associated proteins from background (Table S3) (Chen *et al*, 2013). Tg interactors were similarly analyzed using GO Molecular Function 2018 terms (Tables S3 and 5). The dataset used for the mass spectrometry interactome characterization experiments showing protein identification and quantification are included in Table S8. Spectrum and result files are available via ProteomeXchange with identifier PXD018379.

### Immunoblotting, SDS-PAGE, and Immunoprecipitation

Cell lysates were prepared by lysing in RIPA buffer with protease inhibitor cocktail and protein concentrations were normalized. Lysates were then denatured with 1X Laemmli buffer + 100mM DTT and heated at 95°C for 5 minutes before being separated by SDSPAGE. Samples were transferred onto polyvinylidene difluoride (PVDF) membranes (Millipore) for immunoblotting and blocked in 5% milk in tris-buffered saline, 0.1% Tween-20 (TBS-T). Primary antibodies were incubated either at room temperature for 2 hours, or overnight at 4°C. Membranes were then washed four times with TBS-T and incubated with secondary antibody constituted in 5% non-fat dry milk/TBS-T either at room temperature for 1 hour or overnight at 4°C. Membranes were washed four times with TBS-T and then imaged using a ChemiDoc MP Imaging System (BioRad). Quantification was performed using Image Lab Software (BioRad). For Tg immunoprecipitation, normalized lysates were incubated with M2 anti-flag agarose resin or G1 Anti-DYKDDDDK affinity resin overnight at 4ºC. Resin was then washed four times with RIPA buffer and samples were eluted using 3X Laemmli buffer with 100mM DTT. For immunoblot confirmation of Tg interactors samples were processed exactly as described above for interactome characterization and proteins were eluted once with elution buffer (2% SDS, 1mM EDTA, in PBS) by heating at 95°C for 5 minutes.

### Cycloheximide Chase Assay

In general, 6-well dishes of transfected cells were plated onto poly-D-lysine coated plates and cells were washed twice with 2mL of media treated with cycloheximide (50 μg/mL), then chased with 1mL of cycloheximide-treated media and collected at various time points. Cells were harvested by aspirating media, washing cells twice with 2mL of cold PBS and lysing in 1mL RIPA buffer with protease inhibitor cocktail (Roche, 4693159001). Collected media was spun down at 400x g for 5 minutes to pellet any floating cells. Cell lysate and media was subjected to immunoprecipitation with M2 antiflag agarose resin or G1 Anti-DYKDDDDK affinity resin overnight at 4°C. Resin was then washed four times with RIPA buffer and samples were eluted using 3X Laemmli buffer with 100mM DTT. Eluted samples were separated by SDS-PAGE and transferred to PVDF membrane and probed with primary and secondary antibody as described above.

### ^35^S Pulse Chase Assay

In general, 6-well dishes of transfected cells were plated onto poly-D-lysine coated plates and cells were incubated with methionine and cysteine depleted DMEM supplemented with glutamine, penicillin/streptomycin, and 10% FBS at 37ºC for 30 minutes. Cells were then metabolically labeled in DMEM depleted of methionine and cysteine, and supplemented with EasyTag ^35^S Protein Labeling Mix (Perkin Elmer, NEG772007MC), glutamine, penicillin/streptomycin, and 10% FBS at 37ºC for 30 minutes. Afterward, cells were washed twice with DMEM containing 10 X methionine and cysteine, followed by a burn off period of 30 minutes in normal DMEM. Cells were then chased for the respective time period with normal DMEM, lysed with 500uL of RIPA buffer with protease inhibitor cocktail containing 10mM dithiothreitol (DTT) as described above. Insoluble debris was pelleted by centrifugation at 21,100x g for 15 minutes. Cell lysates were then diluted with 500uL of RIPA buffer with protease inhibitor cocktail and subjected to immunoprecipitation with G1 anti-DYKDDDDK affinity resin overnight at 4°C. After three washes with RIPA buffer protein samples were eluted with 3X Laemmli buffer with 100mM DTT heating at 95°C for 5 minutes. Eluted samples were then separated by SDS-PAGE, gels were dried and exposed on a storage phosphor screen. Radioactive band intensity was then measured using a Typhoon Trio Imager (GE Healthcare) and quantified by densitometry in Image Lab (BioRad).

### EndoH and PNGaseF Treatment

Cells were lysed in either RIPA or TNI (50mM Tris pH 7.5, 250mM NaCl, 1mM EDTA, and 0.5% IGEPAL) buffer with protease inhibitor cocktail, denatured, and digested with EndoH or PNGase F per the manufacturer specifications (New England BioLabs)

