## Supplemental Information for "Thyroglobulin Interactome Profiling Uncovers Molecular Mechanisms of Thyroid Dyshormonogenesis"

### SUPPLEMENTAL FIGURES

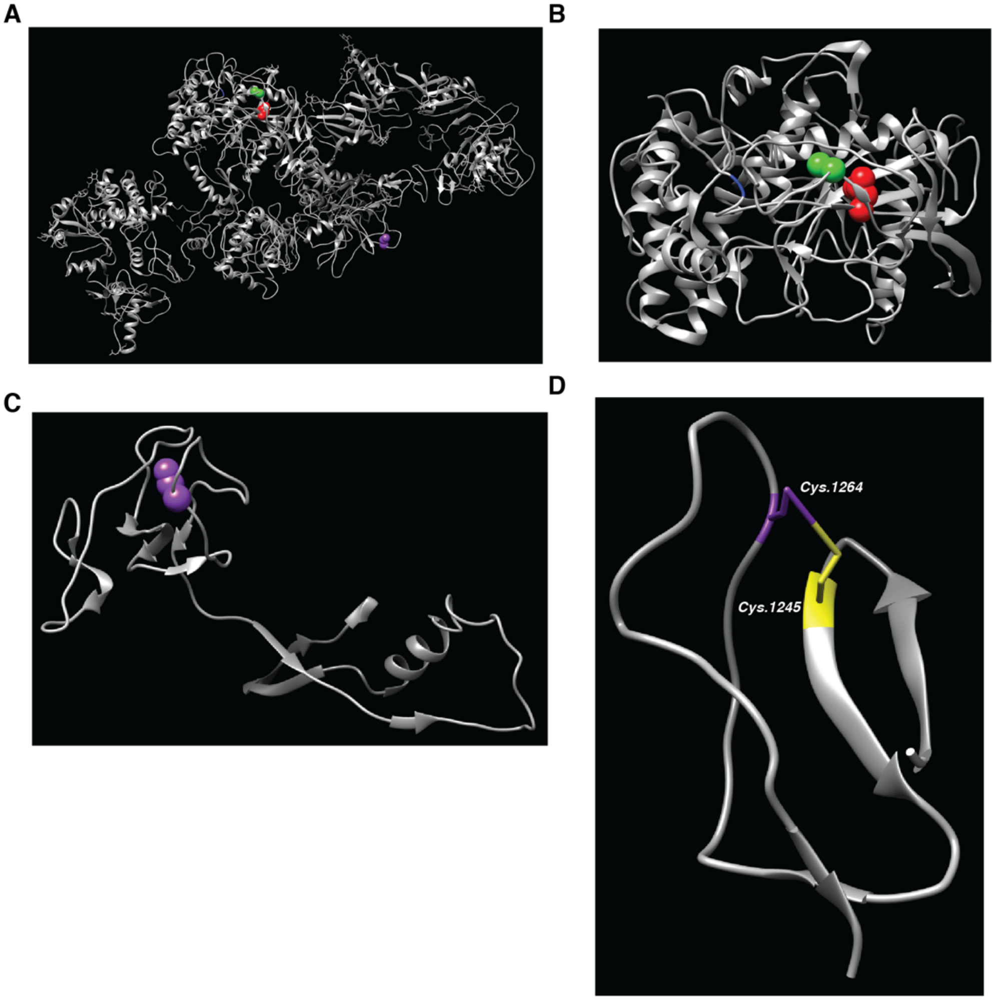


**Figure S1. Structure of thyroglobulin and localization of missense mutations resulting in loss of secretion. A.** Cryo-electron microscopy structure of thyroglobulin (PDB ID: 6SCJ). Only one monomer of the Tg dimer is shown for clarity. Missense mutations resulting in folding-incompetent thyroglobulin are annotated: G2341R (Blue), L2284P (Red), A2234D (Green), and C1264R (Purple). **B.** Close up of the Cholinesterase (ChEL)/CTD domain. ChEL/CTD mutations share close proximity with one another. **C.** Close up of the Hinge/Flap region with C1264R mutation. **D.** Close up of the disulfide bridge formed by C1264 and C1245.

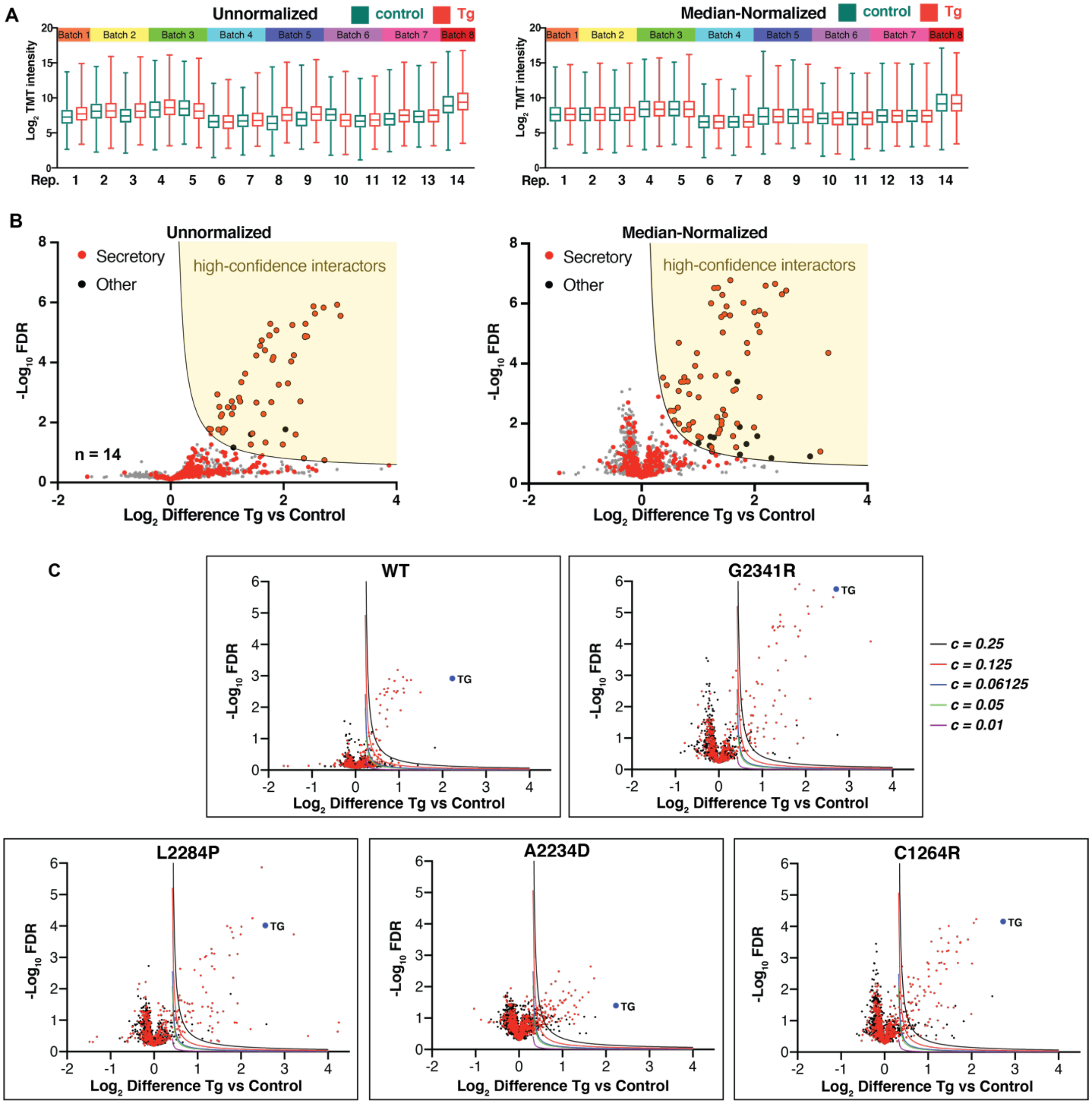
A
Unnormalized
control
Tg
Median-Normalized
Batch 1
Batch 2
Batch 3
Batch 4
Batch 5
Batch 6
Batch 7
Batch 8
Log2 TMT intensity
Rep.
B
Unnormalized
Median-Normalized
Secretory
Other
high-confidence interactors
n = 14
-Log10 FDR
Log2 Difference Tg vs Control
C
WT
G2341R
L2284P
A2234D
C1264R
TG
c = 0.25
c = 0.125
c = 0.06125
c = 0.05
c = 0.01

**Figure S2. Processing of quantitative mass spectrometry data to filter high confidence interactors of Tg. A.** Distribution of TMT reporter ion intensities of all quantified proteins for the Tg (red) and control channels (green). Raw data unnormalized and transformed data based on median-nomalization is shown. TMT channels that were paired in the same mass spectrometry run are indicated by colored bars on top. Source data is shown in Table S1, Sheet 1-2. **B.** Volcano plot for comparison of protein abundances in Tg vs control channels to identify high-confidence interactors using unnormalized TMT intensities or median-normalized TMT intensities. Plots show the average of Tg mutant channels. Median normalization improves the identification of high-confidence thyroglobulin interactors. Source data is shown in Table S1, Sheet 3-4. **C.** Volcano plots displaying the optimization of cut-off parameters for identification of high-confidence interactors for each Tg variant (WT, G2341R, L2284P, A2234D, C1264R). Plots display the average difference in $\log_2$ TMT reporter ions intensities for proteins between the Tg and control channels vs. FDR values (Storey). Cut-offs used to define confident thyroglobulin interactors were optimized as using a previously described method (Keilhauer *et al*, 2015). Cutoffs were designed in such a way that optimized the identification of secretory pathway components compared to non-specific backgrounds proteins, as described in the Materials and Methods section. Shown are specific cutoff lines with different curvature parameters *c*. A cutoff value of $c = 0.05$ was selected to define the confident interactors for each Tg variant. Source data is shown in Table S1, Sheet 5-9. Table S2 shows the list of confident interactors for each variant, which were then combined into a comprehensive list of Tg interactors shown in Table S3 for subsequent analysis.

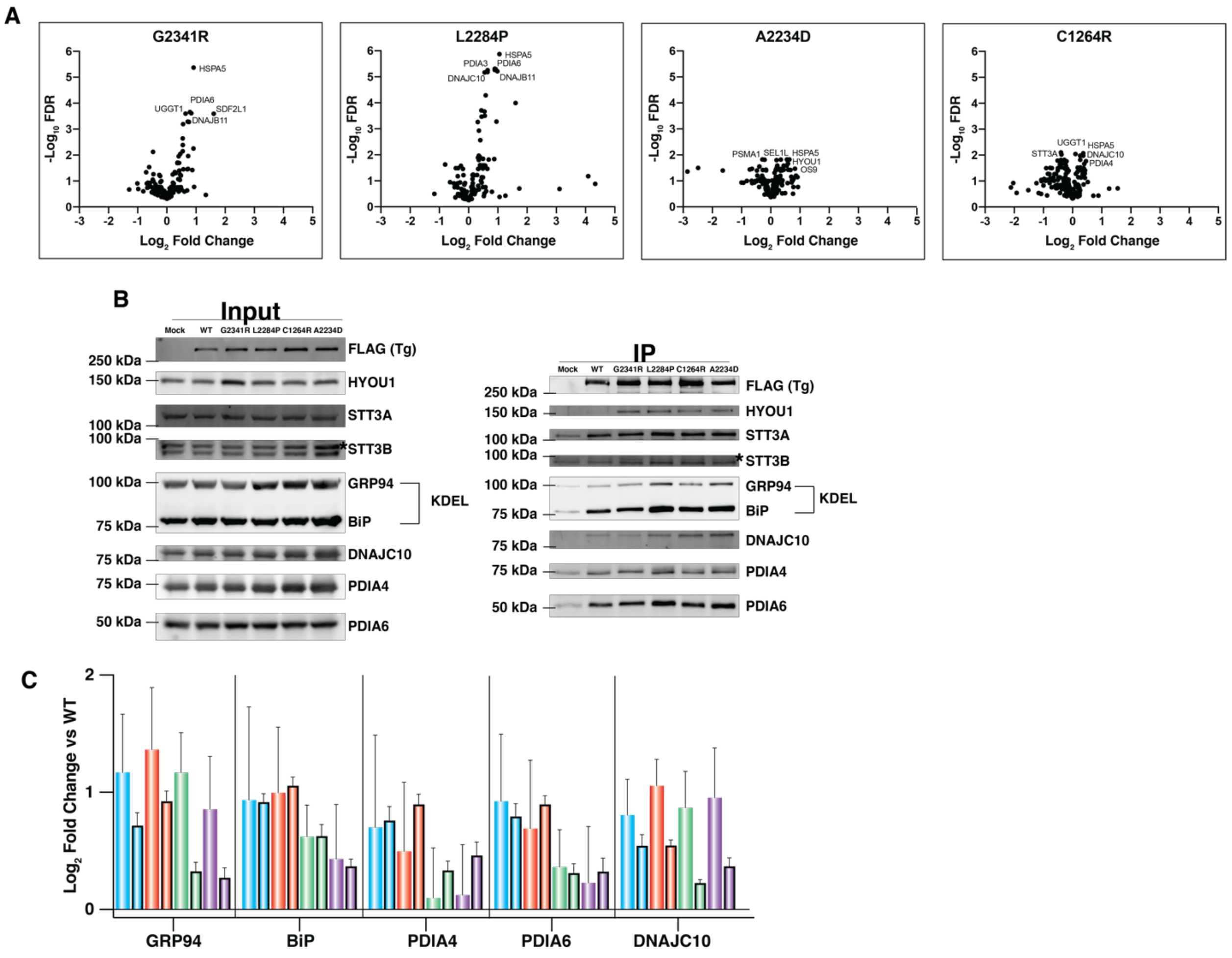


**Figure S3. Comparison of interaction changes between mutant Tg variants and WT Tg. A.** Volcano plot showing the interaction changes of high-confidence interactors between individual mutant Tg variants and WT. Plots display the average $log_2$ difference in TMT reporter ion intensity between the mutant Tg and WT Tg channels for mutant-specific interactors versus the FDR values (Storey) **B.** Representative immunoblots from co-immunoprecipitations confirming select interaction changes between Tg and proteostasis factors. $HEK293^{DAX}$ cells were transiently transfected with the varying Tg constructs or mock as indicated, 0.5mM DSP cross-linker was added to capture transient protein interactions, and co-immunoprecipitations (IP) using anti-FLAG antibody-conjugated beads were carried identical to the Co-AP experiments for the quantitative proteomics experiments. Lysate inputs are shown as controls. Eluted samples were blotted for interaction partners. **C.-D.** Quantification of the Co-IP experiments from **B**. Protein intensities were quantified by densitometry in Image Lab

(Bio-Rad) and protein amounts of interactors in the Co-IP samples were normalized to abundances of Tg variants (FLAG). Error bars show SEM from n = 3 experiments. For comparison, interaction fold changes from TMT-based quantitative AP-MS are shown in bold outlines. Increased interactions with chaperoning and oxidative folding pathways were confirmed.

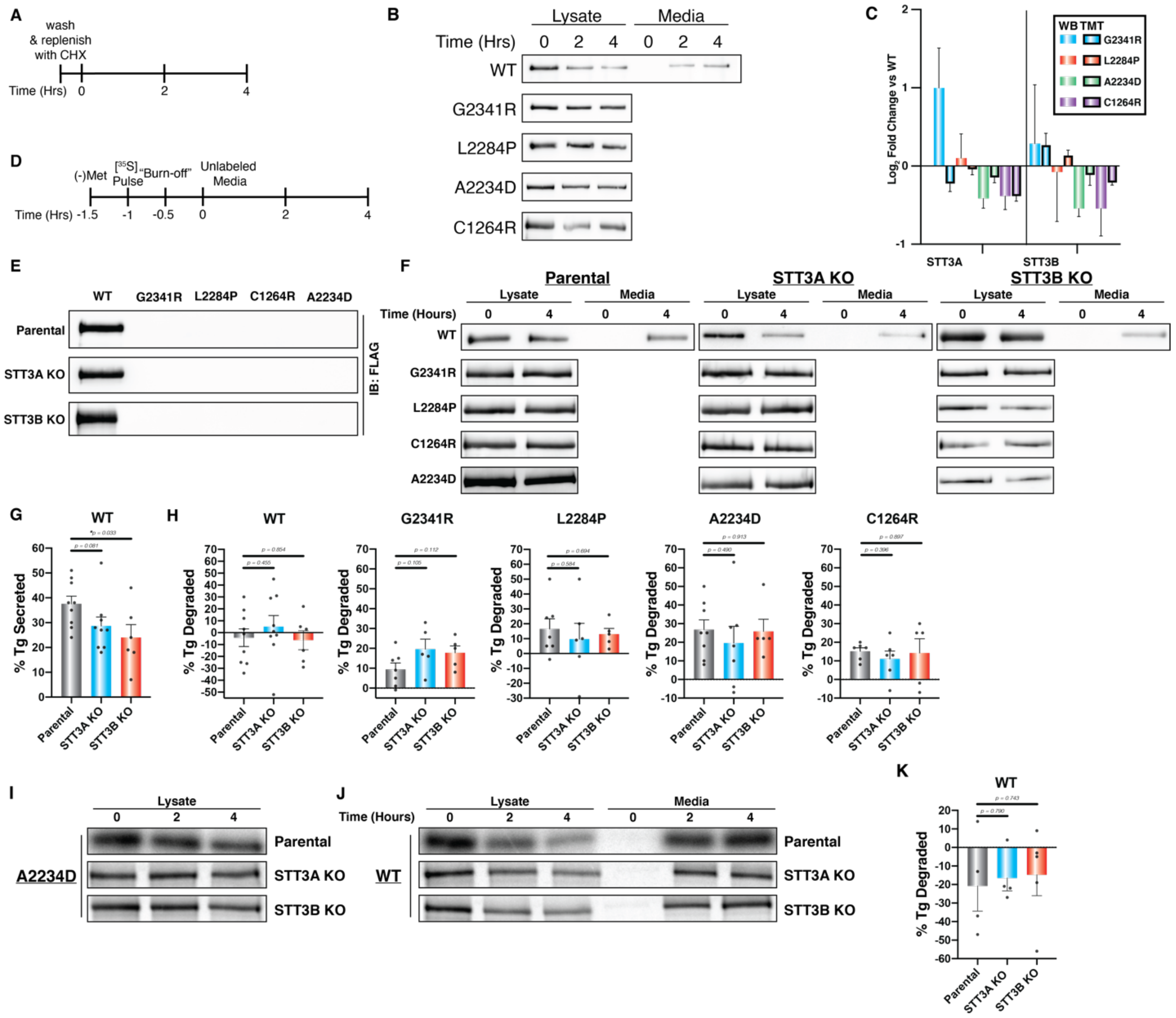


**Figure S4. Cycloheximide chase and pulse-chase assays to measure Tg secretion and degradation rates. A.** Treatment scheme for cycloheximide (CHX) chase assay. **B.** Representative immunoblots from CHX chase assays in Fig. 4E to measure degradation rates of Tg variants. FLAG-tagged Tg variants were transiently transfected into HEK293T cells and treated as outlined in A. Tg from media (WT only) or lysate samples was immunoprecipitated using anti-FLAG antibody beads, resolved by SDS-PAGE followed by immunoblotting. **C.** Confirmation of decreased protein interactions with the OST subunits STT3A and STT3B by Co-IP followed by immunoblots. Error bars shows SEM from n = 3 experiments. For comparison, interaction fold changes from TMT-based quantitative AP-MS are shown in bold outlines. **D**. Treatment scheme for

$^{35}$S-metabolic labeling and pulse-chase assay. **E**. Immunoblots of immunoprecipiated media samples from cells transiently transfected with respective FLAG-tagged Tg plasmid constructs. Neither STT3A or STT3B knockdown impair WT Tg secretion or rescue mutant Tg secretion. **F.** Representative immunoblots from CHX chase assays performed using STT3A and STT3B KO cells to investigate the role of the OST complex isoforms on Tg processing. FLAG-tagged Tg variants were transiently transfected into parental HEK293T or STT3A or STT3B KO cell lines and treated as outlined in A. Tg from media (WT only) or lysate samples was immunoprecipitated using anti-FLAG antibody beads, resolved by SDS-PAGE followed by immunoblotting. Tg protein bands were quantified by densitometry in Image Lab (Bio-Rad). **G.** Plots showing the quantified changes in WT Tg secretion for WT Tg as measured by CHX chase assay in parental, STT3A or STT3B KO cells. % Tg secreted was calculated as $Tg_{media,\ 4h}/\left(Tg_{lysate,\ 0h}+Tg_{media,\ 0h}\right)$. Error bars display SEM of 6-9 biological replicates. Student's parametric t test was used to determine significant changes in Tg secretion and p values are indicated. **H.** Plots showing the quantified changes in Tg degradation as measured by CHX chase assay in parental, STT3A or STT3B KO cells. % Tg degraded was calculated as $1 - (Tg_{lysate,\ 4h}/Tg_{lysate,\ 0h})$ for Tg mutants and $1 - (Tg_{lysate,\ 4h} + Tg_{media,\ 4h})$ for WT. Error bars display SEM of 5-9 biological replicates. Student's parametric t test was used to determine significant changes ($p < 0.05$) in Tg degradation and p values are indicated. **I-J.** Representative autoradiograms of $^{35}$S-pulse chase experiment to measure Tg secretion and degradation. FLAG-tagged Tg variants were transiently transfected into parental HEK293T or STT3A or STT3B KO cell lines and treated as outlined in B. Tg from media (WT only) or lysate samples was immunoprecipitated using anti-FLAG antibody beads, resolved by SDS-PAGE followed by autoradiography. $^{35}$S-labeled Tg protein bands were quantified by densitometry in Image Lab (Bio-Rad). **K.** Plot showing the quantified changes in WT Tg degradation as measured by $^{35}$S-pulse chase experiments. Error bars display SEM of 4-5 biological replicates. Student's parametric t test was used to determine significant ($p < 0.05$) changes in Tg secretion and p values are indicated.

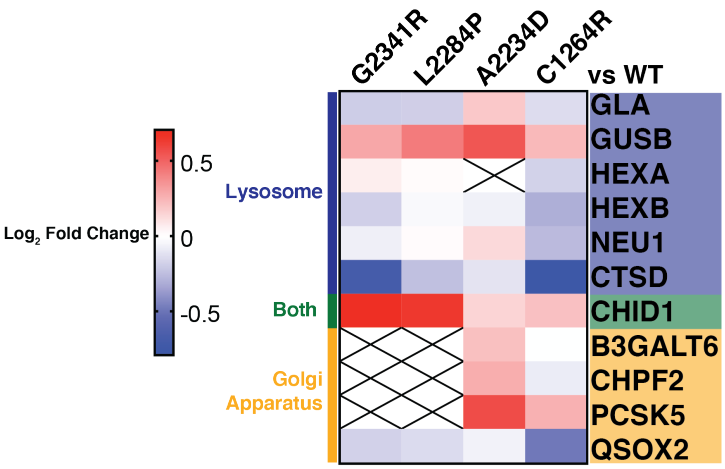


**Figure S5.** Heatmap displaying altered engagement of lysosomal and Golgi related proteostasis components for mutant Tg variants compared to WT Tg.

### SUPPLEMENTAL TABLES

(Tables are included as separate .xlxs or .docx files)

**Table S1.** Source data for Figure S2. Sheets 1 - 4 contain the data used to compare unnormalized vs median normalized TMT intensities to generate plots S2A-B. Median normalization results in better identification of confident Tg interactors. Sheets 3-7 contain the data used to create the plots in S2C. True Tg interactors were further delineated as detailed in S2C and the Materials and Methods section.

**Table S2.** After optimizing normalization methods and identifying optimal cutoffs. Tg interactors were identified for individual constructs.

**Table S3.** Tg interactors found from individual constructs were combined to give a comprehensive list of the Tg interactome. This subsequent list was then used for further data analysis.

**Table S4.** Source data for Figure 2B comparing global Tg vs mock AP TMT intensities expressed as $log_2$ Fold Change (x axis) vs $log_{10}$ FDR (y axis).

**Table S5.** List of Tg interactors sorted based on 2018 Gene Ontology (GO) Biological Process terms

**Table S6.** Source data for Figure 3. $Log_2$ fold change of TMT intensities for interactors comparing CH-associated mutant Tg constructs vs WT Tg. Components sorted by 2018 GO Biological Process terms. For dot plots, individual Tg interactors are plotted together based on GO terms, with each dot corresponding to an individual interactor.

**Table S7.** Table detailing the primers used to perform site-directed mutagenesis on the FLAG tagged WT Tg-pcDNA3.1 plasmid, as described in the Materials and Methods section.

**Table S8.** Raw, unprocessed, protein identification and quantification data from AP-MS experiments used for interactome characterization. Spectra and result files are available via ProteomeXchange under identifier PXD018379.
