## Supplemental Table 7 for "Thyroglobulin Interactome Profiling Uncovers Molecular Mechanisms of Thyroid Dyshormonogenesis"

| Oligo Nucleotide Construct | Sequence | |
| --- | --- | --- |
| G2341R Site-Directed Mutagenesis Forward Primer | | GAGTGGGTGTCTTCCGCTTCCTGAGTTC |
| G2341R Site-Directed Mutagenesis Reverse Primer | | GAACTCAGGAAGCGGAAGACACCCACTC |
| L2284P Site-Directed Mutagenesis Forward Primer | GAAGATTGTTTGTATCCCAATGTGTTCATCCCTC | |
| L2284P Site-Directed Mutagenesis Reverse Primer | GAGGGATGAACACATTGGGATACAAACAATCTTC | |
| A2234D Site-Directed Mutagenesis Forward Primer | AGTTCCATATGaTGCCCCGCCCC | |
| A2234D Site-Directed Mutagenesis Reverse Primer | GAAGCAAGTGGACCAGTTCCTTGG | |
| C1264R Site-Directed Mutagenesis Forward Primer | CAGGGCCATTGATACGTAGCCTGGAGAGC | |
| C1264R Site-Directed Mutagenesis Reverse Primer | GCTCTCCAGGCTACGTATCAATGGCCCTG | |
| D2769X (Untagged WT) Site-Directed Mutagenesis Forward Primer | GCTCTAAGACCTACAGCAAGTGATACAAGGATGACGACGATAAG | |
| D2769X (Untagged WT) Site-Directed Mutagenesis Reverse Primer | CTTATCGTCGTCATCCTTGTATCACTTGCTGTAGGTCTTAGAGC | |
